# The limited place in cognitive space

**DOI:** 10.1101/2022.04.25.489165

**Authors:** Carl J. Hodgetts, Ulrike Hahn

## Abstract

The idea that objects can be represented within multi-dimensional ‘cognitive spaces’ remains popular within psychology and neuroscience, and yet the restrictive topology of such spaces is seldom considered. Here, we show that it is possible, even within a simple set of items, to break such models by imposing neighbourhood relations that are incompatible with a purely spatial representation. These results highlight the fundamental limits of cognitive space as a representational format for complex cognition.

---

How we perceive the similarity between objects in the world is linked intrinsically to how we represent those objects^1,2^. For instance, a cognitive agent that is unable to represent the spatial relationship between different features of an object may easily confuse two objects that share common features but in different arrangements. How one models human similarity, therefore, has significant implications not only for capturing human behaviour across a diverse range of contexts, but for our broader understanding of how human mental representations are structured and organised. Despite this, research across many related disciplines – spanning human and animal psychology, cognitive neuroscience, and machine learning – continues to be dominated by a small subset of models, particularly spatial models of similarity^3^. Here, similarity is broadly conceptualised as distance within a multidimensional cognitive space. According to such a model, an object’s position within the space is determined by its values along a set of relevant psychological dimensions. Proximities between objects or exemplars within this cognitive metric space, therefore, can be considered to reflect similarity, whereby nearby items are highly similar, and distant items are highly dissimilar, as determined by the specific distance metric applied (e.g., Euclidean distance). This theoretical framework has provided the basis for several highly influential models of human recognition within cognitive science^4–7^, and continues to gain popularity within contemporary cognitive neuroscience^8–11^.

Indeed, many of the studies that have applied such spatial models to assess human recognition and conceptual knowledge – both at the behavioural^12^ and neural level^9,10^ – have used stimuli that are well-suited to those models in the first place, in particular objects that vary on a few continuous dimensions. As spatial models operate upon relatively simple object representations (i.e., locations within a *n*-dimensional space), the simple and artificial stimuli often used will not pose serious problems for such models (or individual studies), and thus provide limited insight into the nature or complexity of the underlying object representations – or indeed their neural substrates. For instance, many naturalistic stimuli outside the domain of colours and sounds cannot be easily represented as sets of continuous feature dimensions^13^. Not only are many naturalistic objects characterised by a large number of such dimensions, but these dimensions must be arranged in a particular way to be appropriately categorised, discriminated, and so on^14^. In other words, as soon as representations become structured, knowledge-based, or analogically rich, these basic similarity models appear fundamentally limited^15–17^. Moreover, Euclidean models – and by extension cognitive spaces – place an upper bound on the number of points that can share the same ‘nearest neighbour’^13,18^. Within a 2-D space, for instance, a maximum of 6 distinct items may share the same nearest neighbour, corresponding to the centre and 6 vertices of a regular hexagon. As in the case of structured representation, this may pose few problems for the highly specific and simple stimuli used in many studies, but may lead to systematic distortions within specific stimulus domains, such as those which are hierarchical in nature (e.g., conceptual structures)^13^.

The aim of this paper is to use these basic topological characteristics to demonstrate the limitations of spatial models. Let us first consider the set of items in Figure 1A. These items were generated from a stimulus domain that has been used previously to evaluate structural models of similarity^15,19–22^, allowing us to directly compare spatial models with alternative accounts of similarity – in particular the transformational model of similarity^15,17^. Critically, the transformational model not only assumes the representations of structural information within this domain^15^, but also gives rise to fundamentally different topologies for the similarity space than do spatial models (outlined in Figure 2). The items themselves involve pairs of simple geometric shapes, and these can be described by a basic code where each letter denotes a unique shape (e.g., the comparison ‘square to the left of triangle vs. triangle to the left of square’ can be referred to as *AB/BA*). Under a transformational model, similarities between all possible comparisons can be described via sequences of psychological operators (whereby longer sequences reflect more complex transformations and, in turn, lower perceived similarity). Three psychological operations have been proposed (‘create’, ‘apply’, and ‘swap’), which have been shown to capture similarity judgements and speeded same-different judgements within this domain^15,19,22^ (see Materials and Methods).

**Figure 1.**
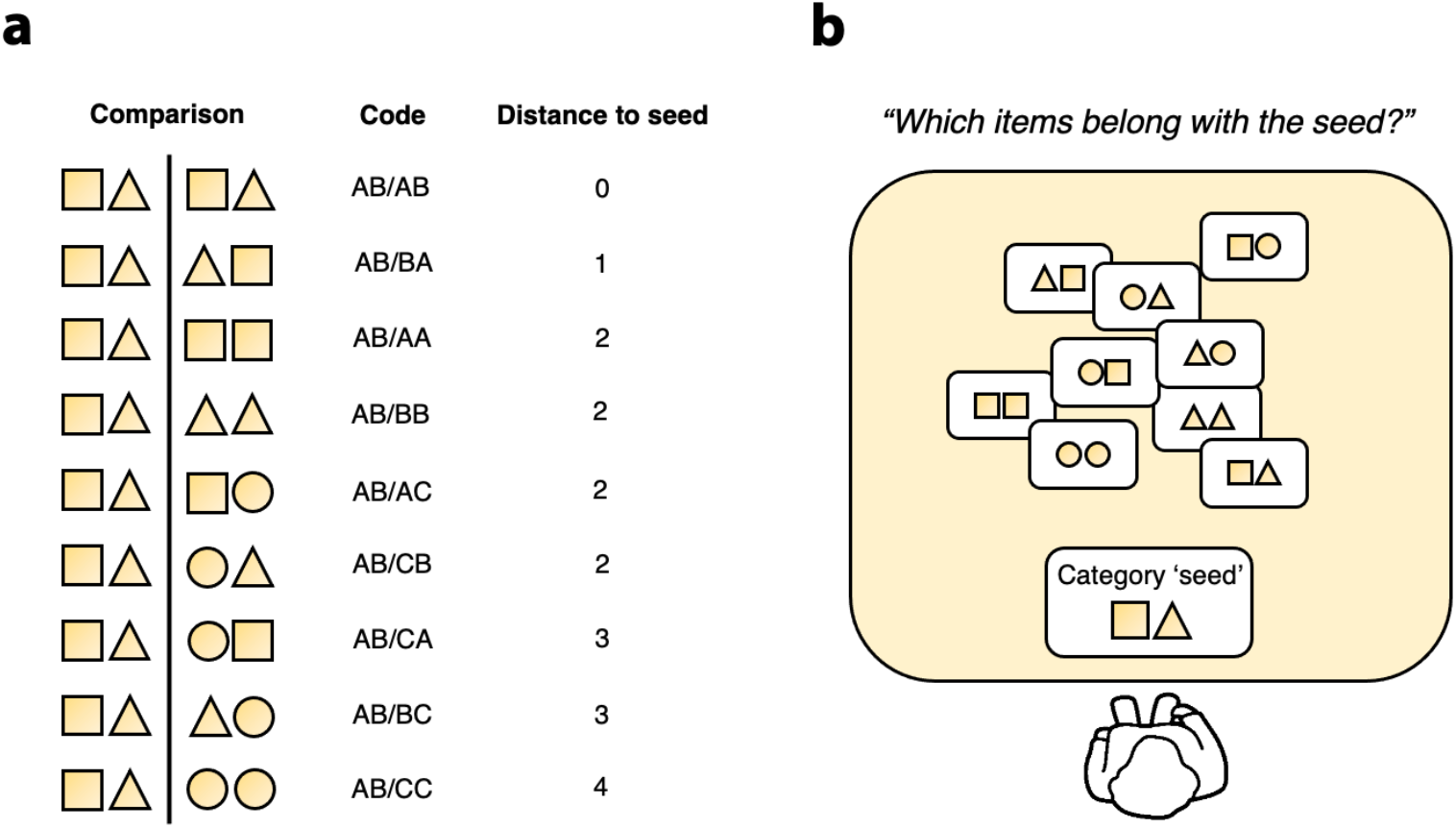
(a) Comparisons between a target item *AB* and nine comparison stimuli. Each item can be described by a code where each feature on the shape dimension can be denoted by a unique letter. The transformational model prediction for each comparison, derived from the coding scheme (see Materials and Methods), is shown. (b) Similarity between such items may be assessed using a free-sorting paradigm. Such a task begins with a ‘seed’ item (item *AB*) and participants must select items that they feel belong with the item.

**Figure 2.**
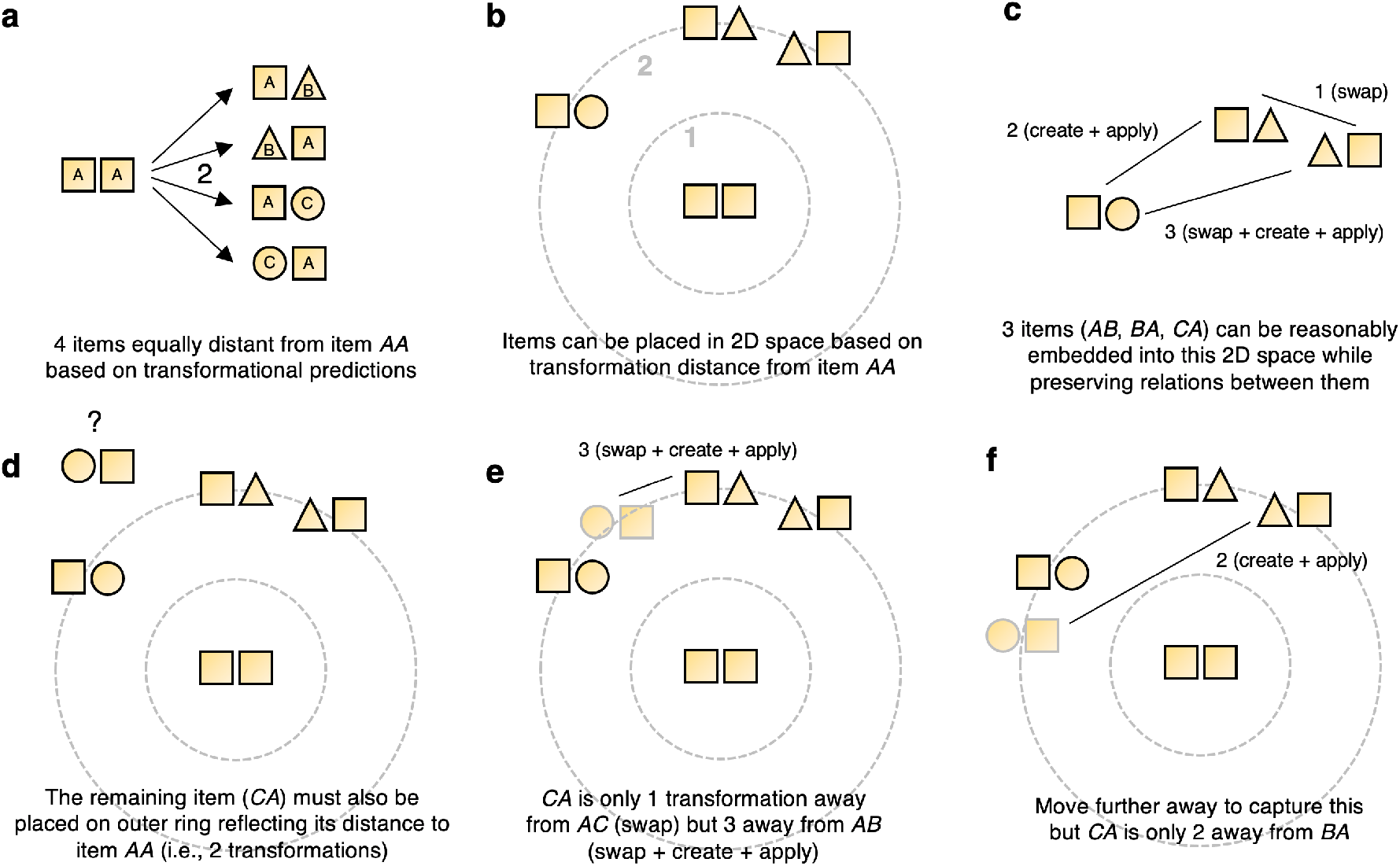
Neighbourhood relations of transformational model distinct from those of a spatial model.

Figure 2 provides an example of how, even for this simple stimulus set, it is possible (under this structural model) to derive neighbourhood relations in this stimulus domain that are incompatible with a purely spatial representation (see Supplementary Figures S1 for a further example). Under the assumed similarity relationships there are four items equally similar to the hypothetical item *AA* (Figure 2A). However, there are no four spatial locations that would allow these items to be equidistant to *AA* whilst also respecting the similarity relations among them. While it is no problem to embed three of these items (see Figure 2B-C), there is no possible location for the fourth item (item *CA*) that respects all relative distances predicted by the transformational model. Crucially, there is nothing complicated about the stimuli, or mysterious about the pairwise transformation distances – both of which are straightforwardly defined^15,19,22^ – it is simply that the more general notion of transformation distance (of which Euclidean or spatial distance is a special case) supports relationships that cannot be embedded within the restrictive topology of spatial distance.

Imagine such similarities being operational within a simple similarity-based task, such as unsupervised categorisation^23^. In such a task, participants are given a specific ‘seed’ item (see Figure 1B) and must decide which items from the broader stimulus set ‘belong’ with this seed item. This type of free-sorting task is used widely because of its close correspondence to category construction in the real-world^24^, and similarity has been shown to be extremely important in determining how items are classified in such a task^25–28^. Past research leads one to expect that the probability of inclusion in the category should decrease as a function of distance to the seed. To show the basic incompatibility between the spatial and transformational model in this domain, one can take the pairwise transformation distances and use multidimensional scaling (MDS) to derive a spatial solution for them. The topological tension of this solution is reflected in a resulting stress value of 0.19, which is considered an unacceptable, or ‘imprecise’, level of data distortion^29^. This tension is also apparent in the annular configuration that is produced, which is commonly seen when fits are poor or when the true underlying configuration is non-annular or incompatible^30^ (Figure 3A, upper panel). Moreover, a correlation plot between these transformation-derived MDS distances and the original transformational distances (Figure 3A, lower panel) shows how specific individual items seemingly ‘break’ the spatial model, in particular items *BB* and *CA* (Figure 2; see also Supplementary Figure 1).

**Figure 3.**
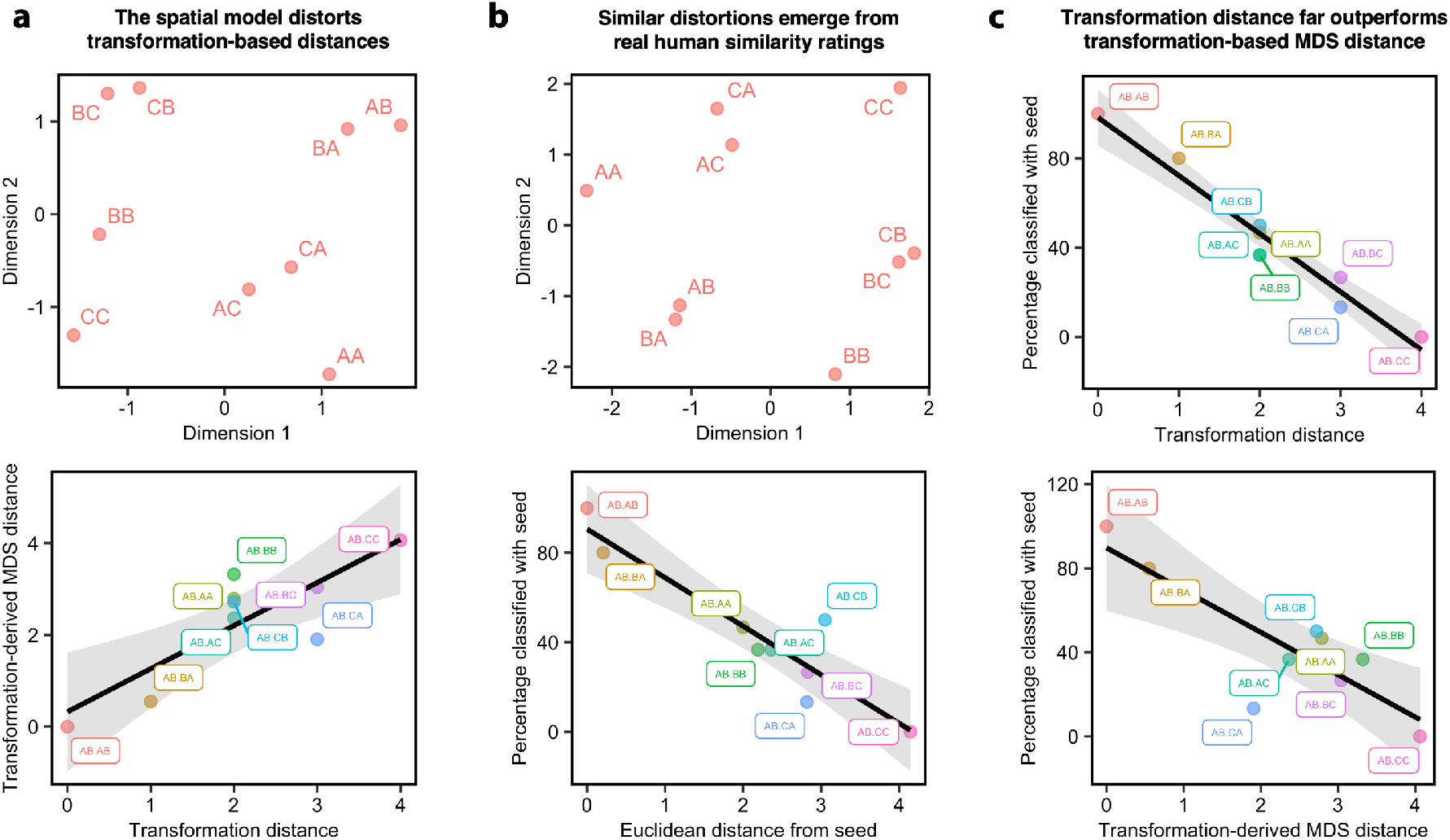
(a) The MDS representation of the pairwise transformational predictions for this item set (top), and the relationship between these distances (labelled, ‘Transformation-derived MDS distances) and the original predictions (bottom). The lower plot shows how MDS distorts the original similarity predictions (see Main text). (b) The MDS representation of the human pairwise similarity ratings for all items in the stimulus set (top), and how these relate to classification performance in the free-sorting task (bottom). (c) The relationship between the transformational model and classification performance (top). This is presented above a plot showing the association between the transformation-derived MDS distances and classification performance. Nine data points are shown on each scatter plot reflecting each item comparison in the task. The 95% confidence interval is also shown.

To demonstrate the psychological relevance of this, we gave this task to 30 participants. This allowed us to confirm the difficulties of the spatial model in capturing the similarity relationships between items in this stimulus domain – both in terms of pairwise similarity judgements and classification behaviour in the sorting task. To derive spatial model predictions for items in the set, we applied MDS to pairwise similarity ratings from an independent set of raters (see Materials and Methods). The distances derived from the MDS representation provided a strong fit of the classification data (r(7) = -0.94, p < 0.001, R^2^ = 0.88; see Figure 3B), yet there are indications that certain item relationships are not easily embedded within this Euclidean solution. Firstly, a fair-to-poor degree of stress is observed for this MDS solution (Stress-1 = 0.11; Guttman, 1965; Kruskal, 1964). And further, the items in this ratings-derived MDS representation also form an annular configuration (Figure 3B), which, under closer inspection, has obvious similarities to the one derived from the transformational predictions. Finally, while the spatial model provides a good fit of the classification data, the transformational model’s fit is near-perfect (r(7) = -0.98, p < 0.0001), accounting for 95% of the variance (Figure 3C). A likelihood ratio based on these fits suggest that the data are 51 times more likely to occur under a transformational model (see ref. ^31^).

Two final considerations further confirm the topological difficulties of the spatial model. First, we used the transformation-derived MDS distances (depicted in Figure 3A) to predict classification behaviour in our human participants. Although statistically significant (r(7) = -0.84, p = 0.004, R^2^ = 0.71), the fit is less accurate than that obtained from original transformational model predictions, implying that MDS distorts these predicted similarities (Figure 3C). This is confirmed by an increased Stress-1 value of 0.19. Second, the residuals that were problematic when deriving an MDS representation of the transformation distances (see Figure 3A), are similarly problematic when those distances are derived from human similarity judgements (namely, items *BB* and *CA*). Notably, the original transformational predictions fit these comparisons very well (see Figure 3C, upper panel) suggesting that these poor fits relate to the spatial model in general and not, for example, the sample-size from which the pairwise ratings were derived.

Multidimensional cognitive spaces, in which similarity is determined by Euclidean distances between objects, are central to many influential models of human cognition^4–7^, and have recently been proposed as the “primary representational format” for information coding in the brain^10^. However, are such models even appropriate for estimating similarity between simple stimuli when they are not amenable to those models in the first place? Here, we showed how a stimulus domain that has been used successfully to provide of evidence for transformational models of similarity in the context of similarity ratings^15,22^ and speeded same-different judgements^19^, can be used to create similarity relationships that cannot be simply embedded within a Euclidean metric space (Figure 2). Indeed, one of the fundamental limitations of so-called ‘spatial codes’ is that it places an upper bound on the number of distinct items that can be equally similar to any other^13^. While this may pose few problems within specific and artificial stimulus domains that are designed specifically with Euclidean spaces in mind (e.g., manipulating continuously valued dimensions), it places a fundamental limit on such representations as a generalised model of human cognition, whereby multiple, complex object and events must be co-represented, remembered and discriminated.

Our approach revealed that spatial models can systematically distort the relationship between objects, even within a relatively simple stimulus domain. Such distortions were evident when spatial/MDS representations were derived from human similarity ratings (and then applied to novel classification data in a free-sorting task) and reflected those seen when spatial representations were derived directly from the predictions of the transformational model. This analysis demonstrated how certain item similarities were simply not compatible, leading to a lack of precision in model fits – even within this relatively ‘simple’ stimulus domain.

Thus, while such spatial models may be useful in highly specific contexts, these results strongly highlight their fundamentally limits as a representational format for complex human cognition.

## Materials and methods

### Participants

A total of 30 Cardiff University students completed the spontaneous categorisation task (mean age = 19.2, range = 18-25). An independent sample of 7 participants also completed a pairwise similarity rating task. For both tasks, participants were allocated course credit for taking part and were tested individually in a cognitive testing laboratory. This study was approved by the School of Psychology local ethics committee, and all participants provided informed consent prior to taking part.

### Stimulus set

The stimulus set consisted of nine pairs of geometric shapes (Figure 1). The seed item in the free-sorting task (see below) comprised a square to the left of a triangle (represented by the code *AB*). Note, we will refer to each potential item (a shape pair) and comparison (a comparison between two shape pairs) using a letter code, where every letter corresponds to a unique feature on the shape stimulus dimension (*A* = square; *B* = triangle; *C* = circle). For instance, the comparison between the ‘seed’ item and the two circles can be labelled *AB/CC* (see Figure 1). As there was a single category reference (the category seed), the only relevant comparisons for modelling the data were those between task items and the category seed, that is, not all pairwise comparisons are carried out for this specific task.

### Tasks and procedure

#### Free sorting task

In this task, participants were given initially the category ‘seed’ (item *AB*, see ‘Stimulus set’). A ‘seed’ item is a single stimulus that acts as a category reference^23^. After studying this ‘seed’, the eight comparison items were presented and the whole array was available (see Figure 1B). Participants were instructed to use the ‘seed’ as a basis for their judgements and to choose items that they felt belonged with it. The seed method was applied because engaging participants in all possible pairwise comparisons between individual items would have resulted in several ‘repeats’, that is, equivalent comparisons with matching predictions. Having participants compare the seed with all other task items ensured that each comparison was unique and could be easily mapped onto the model predictions (see ‘Model predictions’ below). To make the task more engaging, we presented each item as runes on an alien spacecraft and created a back-story in which the participants were required to classify spacecrafts as hostile or friendly. The back-story was presented on an A4 sheet in front of the participant. At the bottom of the sheet was an envelope that concealed category seed. The spaceships were 8cm x 5cm with the pair located in the centre of each. Each shape was 0.7cm x 0.7cm and was separated by a horizontal distance of 0.2cm. Stimuli were presented on laminated card that was 12cm x 6cm. Participants were tested individually and were first instructed to read the back-story before inspecting the category seed. After reading the story they were asked to open the envelope and reveal the category seed. After inspecting the seed item, they were handed the nine task items and asked to organise them as described on the instruction sheet. After completing the sorting task, participants were asked how they classified the items.

#### Similarity rating task

A separate similarity rating task was used to derive pairwise similarities for MDS (see ‘Model predictions’ below). The task itself was presented on a 19” LCD monitor using Inquisit software (Millisecond Software, Seattle, WA). There were 81 trials in total (9 × 9) and each trial involved a comparison between two items from the stimulus set, one presented on the left of the display, and one on the right. Each possible comparison was presented twice, counterbalancing for the left-right position of items. As in the classification task, each item was a pair of geometric shapes (Figure 3) superimposed on a line drawing of a spacecraft. Participants could take as long as they wanted on each trial. The task took approximately 7 minutes to complete.

### Model predictions

#### Spatial model

According to the spatial model, an object’s position within the space is determined by its values along a set of relevant psychological dimensions. Distance between exemplars within this psychological metric space, therefore, can be considered to reflect similarity, whereby nearby items are highly similar, and distant items are highly dissimilar. Distance in geometric models is defined as:

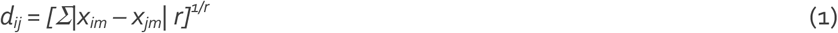

where *d*_*ij*_ is the distance between objects *i* and *j, x*_*im*_ is the value of object *i* on dimension *m* and the parameter *r* refers to the spatial metric employed (i.e., Euclidean [r = 2] or city-block [r = 1]). In addition to being a model of similarity and mental representation, the spatial model also provides a technique for summarising and displaying similarity data. The statistical procedure of multidimensional scaling (MDS)^3^ will generate, as its output, a spatial representation of minimum dimensionality that preserves the proximities in the input data as best possible. The goodness of this fit is reflected in the ‘stress’ of a given solution (known as Stress-1). A high ‘stress’ value indicates that the input data has been distorted greatly in generating the output. The distances between objects in this output representation can be used to predict patterns of data in appropriate psychological models or fit related sets of similarity data^4,32^. To derive spatial model predictions, we applied MDS to pairwise similarity ratings from an independent set of raters using ‘cmdscale’ in R (see above). Input data corresponded to a 9 × 9 similarity matrix with each cell corresponding to the mean similarity rating for that comparison (averaged across N = 7). Similarity predictions between each item and the seed stimulus were derived by calculating the Euclidean distances between each item and the seed.

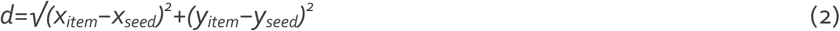

### Transformational model

The similarity predictions for the transformational model were derived from a coding scheme described previously^15^. This scheme specifies three operations that can be applied to change the representation of one stimulus in this set into another. Each operation takes a reference object and modifies it, as follows: 1) *‘Create’*– taking the reference object we apply this operation to create a new feature that is unique to the target pair. For example, if the base pair consists of two squares, and the target pair consists of a square and a circle, we have to create the feature ‘circle’ as this is not present in the base pair; 2) *‘Apply’* - this operation takes an object or feature that is currently available (either via its presence in the base stimulus or via the ‘Create’ operation) and applies it to one or both of the objects in the target item. Here, applying a shape that is readily available (on a representational level) to both shapes in the target is equivalently difficult to applying it to one. Thus, once the ‘circle’ feature has been created (using operation 1), it is considered equivalently effortful to apply it to one, or both, of the square objects; 3) *‘Swap’* – this swaps the relative spatial position of individual features or whole objects. These transformations are applied bi-directionally to each stimulus comparison, and the maximum transformation distance is taken as the prediction for that comparison^17,19^.

### Transparency and openness

Task materials (e.g., stimuli and run files), data (e.g., key performance metrics across tasks and species), and analysis scripts/code to run analyses/generate figures have been made available via the Open Science Framework and can be accessed at the following peer review link: https://osf.io/346pv/. Any questions and additional requests for data can be sent to the corresponding author via email.

## CRediT author statement

**Carl J. Hodgetts:** Conceptualization, Investigation, Formal Analysis, Data Curation, Visualization, Writing – Original Draft, Writing - Reviewing and Editing. **Ulrike Hahn:** Conceptualization, Investigation, Formal Analysis, Writing – Original Draft, Writing - Reviewing and Editing.

## Acknowledgements

This work was supported by the Biotechnology and Biological Sciences Research Council [BB/V010549/1] and the European Commission New and Emerging Science and Technology (NEST) Scheme [516542].

## Notes

https://osf.io/346pv/

### Competing Interest Statement

The authors have declared no competing interest.

