## Supplementary Information for "The limited place in cognitive space"

### Supplementary Figure 1

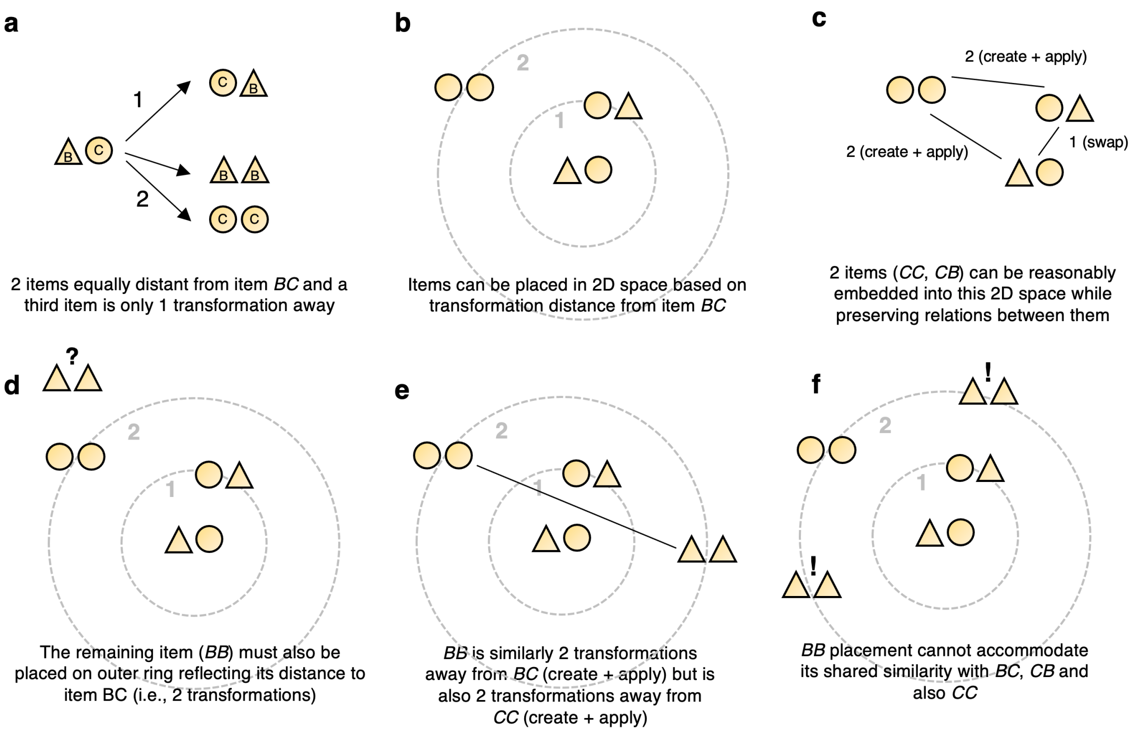


**Supplementary Figure 1.** A further example of how neighbourhood relations of transformational model are distinct from those of spatial models (see Main text)
